# aiCRISPRL: An Artificial Intelligence Platform for Stem Cell and Organoid Simulation with Extensive Gene Editing Capabilities

**DOI:** 10.1101/2022.06.18.496679

**Authors:** WR Danter

## Abstract

CRISPR-Cas9 (clustered regularly interspaced short palindromic repeats/CRISPR-associated nuclease 9) provides powerful gene-editing tools that are applicable for gene therapy of a variety of diseases including, but not limited to cancer, rare diseases, and heart disease. In the current study, we first re-examined our artificial stem cell and organoid simulations that were generated by our literature validated DeepNEU AI platform from the perspective of gene-editing. We then evaluated the aiCRISPRL (aiCRISPR-Like) application of the DeepNEU platform by directly comparing the CRISPR-Cas9 gene-editing approach with the DeepNEU derived aiCRISPRL capabilities using artificial simulated HeLa cells (aiHeLa). To accomplish this, we evaluated the aiCRISPRL like capabilities of DeepNEU to introduce a series of specific mutations into the MutS homolog 2 (MSH2) gene to assess DNA Mismatch Repair (MMR). This approach permits a comparative assessment of CRISPR-Cas9 and aiCRISPRL technologies following the introduction of specific MSH2 mutations. When combined with our previous body of gene editing research, the current data indicates that aiCRISPRL is an advanced AI platform technology that can be used for rapid prototyping and multiple scenario simulation in genomic research to complement wet-lab based gene-editing technologies.

## INTRODUCTION

In the last decade, gene editing technologies like CRISPR-Cas9 and TALEN have revolutionized our ability to rewrite the genetic code, offering unprecedented opportunities to advance human health through disease modeling, therapeutic interventions, and drug discovery. Central to this technological revolution is CRISPR-Cas9, which, through its precise and efficient DNA modification capabilities, has been pivotal in enhancing our understanding of genetic diseases while fostering the development of an increasing array of novel treatments [1,2]. CRISPR technology’s broad implications and innovative applications extend well beyond traditional gene editing, opening new avenues in genomic research and medicine [3]. However, despite its transformative potential, CRISPR-Cas9 encounters significant challenges, including off-target effects and ethical concerns that currently limit its broader application [4].

Parallel to these advancements in vitro, artificial intelligence (AI) is playing an increasingly critical role in the life sciences, offering innovative solutions that complement and enhance traditional research methodologies. At the forefront of this integration is DeepNEU©, an advanced AI/ML platform that simulates pluripotent stem cells [5], human organoids [6], and the aiHumanoid [7] to unravel the complex interplay of genetics in disease, without the limitations associated with direct genetic manipulation. This synergy between AI and gene editing technologies heralds a new wave of precision medicine, where the predictive power of AI and the editing precision of CRISPR could together significantly reduce discovery and therapeutic development timelines [8].

In this study, we present an innovative approach that combines the strengths of CRISPR-Cas9 gene editing with the predictive capabilities of the DeepNEU platform. Our work introduces the aiCRISPRL application, a novel AI-driven model that simulates CRISPR-like gene editing outcomes with remarkable accuracy. This approach not only navigates the ethical and practical challenges associated with direct genetic modifications but also offers a scalable and cost-effective pathway for exploring genetic interventions through computer simulations.

By directly comparing the outcomes of CRISPR-edited [9] and AI-simulated aiHeLa cells with MSH2 mutations, our research highlights the complementary nature of these technologies. This comparative analysis provides validation for aiCRISPRL’s predictive accuracy while demonstrating its potential to refine gene editing strategies, optimize therapeutic targets, and improve our understanding of gene repair mechanisms.

This research represents a significant advance in the fields of genetic research and pharmacogenomics, bridging the gap between the physical and computational realms of science. It underscores the potential of integrating AI with gene editing technologies to not only expedite the pace of genetic research but also to address the ethical and practical challenges currently facing the field. Our study sets a new standard for interdisciplinary research, offering insights that could lead to a paradigm shift in how genetic research is conducted and applied in medicine.

## METHODS

The DeepNEU platform is an advanced literature-validated unsupervised machine learning platform [5-7,10-21]. Here we are presenting the updated database of DeepNEU (v8.4). this version contains the previous database (v8.3), in addition to new genotypic and phenotypic concepts and new pathway relationships.

The goal of this project was to generate simulated HeLa cells (aiHeLa) by applying aiCRISPRL gene editing of the simulated version of induced pluripotent stem cells (aiPSC) genome to introduce consensus gene mutations typical of HeLa cells [22-26].

Briefly, the values of the high dimensional input or initial state vector (N=7529 concepts) of the aiPSC model, were all set to zeros except for the four transcription factors OCT3/4, SOX2, KLF4 and cMYC. These four factors were given a value of +1 indicating that they were turned on for the first iteration and then allowed to evolve over successive iterations until a new system wide steady state is achieved. The detailed methodology for generating aiPSC simulations has been reported in detail previously [5-7,10-21].

Previous versions of the DeepNEU platform have successfully simulated a broad range of aiPSC derived cell types including neural stem cells, cardiac myocytes, skeletal muscle cells, lung cells, brain cells, ovarian cancer cells, lung adenocarcinoma cells, and natural killer cells (NK) [5-7,10-21]. Using a similar approach and a consensus gene mutational profile from multiple sources [22-26], we used the gene-editing capabilities of aiCRISPRL to create simulations (aiHeLa) that are most like original HeLa cells. Importantly, except for the simulated HPV18 infection, the aiHeLa cells were created to be devoid of all external contaminations that plague HeLa variant clones that have emerged since 1951.

### Simulation of HeLa cells (aiHeLa)

Once the aiPSCs were validated against the current literature they were transformed by aiCRISPRL into aiHeLa cells using a gene mutational profile derived from the peer-reviewed literature [22-26]. This aiHeLa mutation profile included GOF mutations in CD24, CD44, CD95(FasR), HPV-E2, 5, 6, and 7. Aneuploidy, an important feature of HeLa cells, was also locked ON and the cells were maintained in simulated DMEM/F12 media supplemented with the addition of doxycycline. This information is summarized in Table 1 below.

**Table 1:** Summary of aiPSC and aiHeLa cell simulation protocol

|  |  |  |
| --- | --- | --- |
| Concept | Recipe | Component/Edits |
| aiPSC | Yamanaka transcription factors | OCT4, cMYC, KLF4 and SOX2 – GOF gene edits |
| aiHeLa | Mutational profile | CD24, CD44, CD95(FasR), HPV-E2,5,6,7 – GOF gene edits, Aneuploidy turned ON |
| DMEM/F12 media | Basic media | [eNa+], [ePyruvate], [eZinc], Choline, FGF2/bFGF, Folic Acid, HEPES, HSA/Albumin, Hypoxanthine, Inositol, KSR, Nicotinamide, ROCK1/2 inhibitor, VitB12/Cobalamin, VitB6/Pyridoxine |
| DMEM/F12 media | Supplements | Doxycycline turned ON |
| Environment | Settings | [O2]=21%, [CO2]=5%, Temperature=37° C - locked ON |
| Patient factors | Age, Smoking | 31 year old smoker – Locked ON |

Outcome predictions from the HeLa cell simulations (aiHeLa) were directly compared with published data regarding markers for HeLa cells [22-26]. Raw output data were transformed to the 0 to +1 scale using the formula y=(x+1)/2. Final expression values > 0.1 were considered as expressed or upregulated for genes/proteins or present for phenotypic features while values ≤0.1 were considered as downregulated, not expressed, or absent.

To evaluate the ability of the DeepNEU platform to simulate HeLa cells, a consensus feature profile was created from the published literature [22-26]. Each feature was identified in at least two published references. The final HeLa cell feature profile (N=10) includes ALDH1, Aneuploidy, CD24, CD44, CD95(FasR), HPV-E2, HPV-E6, HPV-E7, and Telomerase in the setting of chronic HPV-18 infection. The accuracy of simulation predictions was compared to the published literature using the unbiased binomial test. A test p-value <0.05 was used to reject the null hypothesis (H^0^) that the simulated HeLa cell (aiHeLa) profile could not reproduce the documented wet-lab results.

### aiCRISPRL-simulated gene editing of the MSH2 gene in aiHeLa cells

Following the successful creation and literature validation of the aiHeLa simulations, we turned our attention to modifying the MSH2 gene to produce a range of LOF mutations. To allow a direct comparison with published data we followed the approach and analysis presented in [27]. These authors compared wildtype MSH2 to seven specific MSH2 LOF mutations in HeLa cells that were created by CRISPR gene editing. Our modified approach is summarized in Table 2.

**Table 2:** Summary of aiCRISPRL gene editing of MSH2 conducted in simulated HeLa cells (aiHeLa)

| Simulation Name | aiCRISPRL Gene Editing | MSH2-% (Scaled*) |
| --- | --- | --- |
| aiHeLa | None (WT) | 100 (+1) |
| aiHeLa-MSH2-KO | MSH2-deleted | 0 (-1) |
| aiHeLa-RD03 | MSH2 G674R1 | 11 (-0.780) |
| aiHeLa-RH07 | MSH2 G674R2 | 5.2 (-0.896) |
| aiHeLa-DA02 | MSH2 G674D1 | 4.8 (-0.904) |
| aiHeLa-DC08 | MSH2 G674D2 | 3.4 (-0.932) |
| aiHeLa-AG11 | MSH2 G674A1 | 30 (-0.400) |
| aiHeLa-AH07 | MSH2 G674A2 | 35 (-0.300) |

Legend: (Scaled*) - the % MSH protein expression was converted to the -1 to +1 range used as inputs to the DeepNEU/aiCRISPRL platform.

Specific MSH2 LOF mutations were assigned a simulation name and programmed to reproduce the associated changes in the amount of MSH2 protein expression as per Table 2.

### Loss of Function (LOF) Scores

In their paper [27] the authors defined and validated a LOF score for a group of seven CRISPR edited MSH2 mutations plus WT that they evaluated. To develop this score, the HeLa cells were treated with 6-Thioguanine (6-TG). Their MOA-based reasoning led them to conclude that the cells with intact MMR pathways should be sensitive to 6-TG exposure while the cells with defective pathways (dMMR) should show varying degrees of resistance as measured by cell death. Their LOF score was calculated from the Log (base 2) of the ratio of 6-TG to placebo treatment effects in each of the gene-edited and WT cells. Positive LOF scores imply deleterious mutations while negative scores suggest more neutral mutations. Analysis of their data revealed a Pearson correlation r of 0.770 between the severity of the MSH2 mutations and HeLa cell death.

Our aiCRISPRL generated LOF scores were modified in that the amount of HeLa cell apoptosis was substituted for MSH2 mutation severity since 6-TG primarily induces cell death through apoptosis [28]. In addition, the DeepNEU input scaling range from -1 to +1 results in some negative values for aiHeLa cell apoptosis necessitating a reversal of numerator and denominator to derive the ratio used to produce the LOF score. As in [27], positive LOF scores imply deleterious mutations, and negative scores indicate typically more neutral mutations. In the present project the correlation between the LOF score and MSH2 protein expression was evaluated by calculating the Pearson correlation coefficient (r).

## RESULTS

### The Updated DeepNEU database

In the present study, v8.4 (2024) of the DeepNEU database, which is characterized by modest upgrades to its previous version 8.3, was used [18]. Specifically, v8.3 had 7447 concepts and 69581 nonzero causal relationships. Version 8.4 has 7529 concepts and 70587 relationships. This means that for every nonzero concept in the causal relationship matrix, there are approximately 9.4 incoming and outgoing causal relationships.

The previously implemented early stopping algorithm (v8.3) was retained in this updated version. For the purposes of this study, we chose to use the results after 18 iterations, as determined through a three valued moving average, which minimized any evidence of overfitting. In addition, a detailed review of the relationships revealed that the pretest probability of a positive outcome is 0.660 and the probability of a negative prediction is therefore 0.340. These values were used to optimize the binomial test by eliminating system biases prior to its use.

### Previous gene editing examples with CRISPRL

On review of our previous body of work [5-7,10-21], the aiCRISPRL functions of the DeepNEU platform successfully used gene editing to create single-gene mutations, multiple mutations and even edit the entire genome of the SARS-CoV-2 and NIPAH viruses. These edits created both loss of function (LOF) and gain of function (GOF) mutations. A summary of these data from previously published DeepNEU research projects from the perspective of successful gene editing is presented in Table 3 below.

**Table 3:** Examples of aiCRISPRL-Like gene-editing applications of the DeepNEU platform

| Date | aiStem Cell | aiNSC | aiSkMC | aiLUNG | aiOrganoid | Disorder | LOF Mutations | GOF Mutations | Reference |
| --- | --- | --- | --- | --- | --- | --- | --- | --- | --- |
| 2019 | Yes | Yes | No | No | No | RETT Syndrome | MeCP2 | None | 5 |
| 2019 | Yes | No | Yes | No | No | IOPD | GAA | None | 11 |
| 2020 | Yes | No | No | No | No | SARS-CoV-2 #1 | Whole genome | Whole genome | 12 |
| 2021 | Yes | No | No | No | Whole Brain | Alzheimer's Disease | None | APO-E | 6 |
| 2021 | Yes | No | No | No | Lung | SARS-CoV-2 #2 | Whole genome | Whole genome | 13 |
| 2021 | Yes | No | No | No | Lung | LUAD | TP53 | EML4-ALK fusion, Amplicon 11q13, NFKBIA, NKX2-1 | 14 |
| 2021 | Yes | No | No | Yes | No | SARS-CoV-2 #3 | Whole genome | Whole genome | 15 |
| 2021 | Yes | No | No | Yes | Whole Brain | MLD | ARSA | None | 16 |
| 2021 | Yes | No | No | No | Ovary | HGSOC | Many | Many | 10 |
| 2022 | Yes | No | No | No | Whole Brain | NIPAH encephalitis | Whole genome | Whole genome | 17 |
| 2023 | Yes | No | No | No | 21 Organoids | Gram neg Sepsis | None | cMYC, KLF4, SOX2, OCT4 | 7 |
| 2023 | Yes | No | No | No | 21 organoids | SCD | Beta globulin | None | 18 |
| 2023 | Yes | No | No | No | 21 organoids | PAC | P53, CDKN2A | KRAS, SMAD4 | 19 |
| 2024 | Yes | No | No | No | 21<br>organoids | AD | None | APOEs,<br>APP,<br>PSEN1 and<br>PSEN2 | 20,21 |

### The aiCRISPRL-derived aiHeLa Cells Simulations – LOF Score Analysis

The aiCRISPRL gene modifications to the aiPSC described above resulted in aiHeLa cells with a profile consistent with the literature-derived ten feature HeLa profile including ALDH1, Aneuploidy, CD24, CD44, CD95, HPV-E2, HPV-E6, HPV-E7, and Telomerase in the setting of chronic HPV-18 infection..

The probability that all ten HeLa markers were expressed by chance alone is 0.016 based on the unbiased binomial test. The Null hypothesis (H^0^) can therefore be rejected in favor of the alternate hypothesis (H^1^) that the simulated HeLa cell (aiHeLa) profile can accurately reproduce published wet lab results. These results are summarized below in Figure 1.

**Figure 1:**
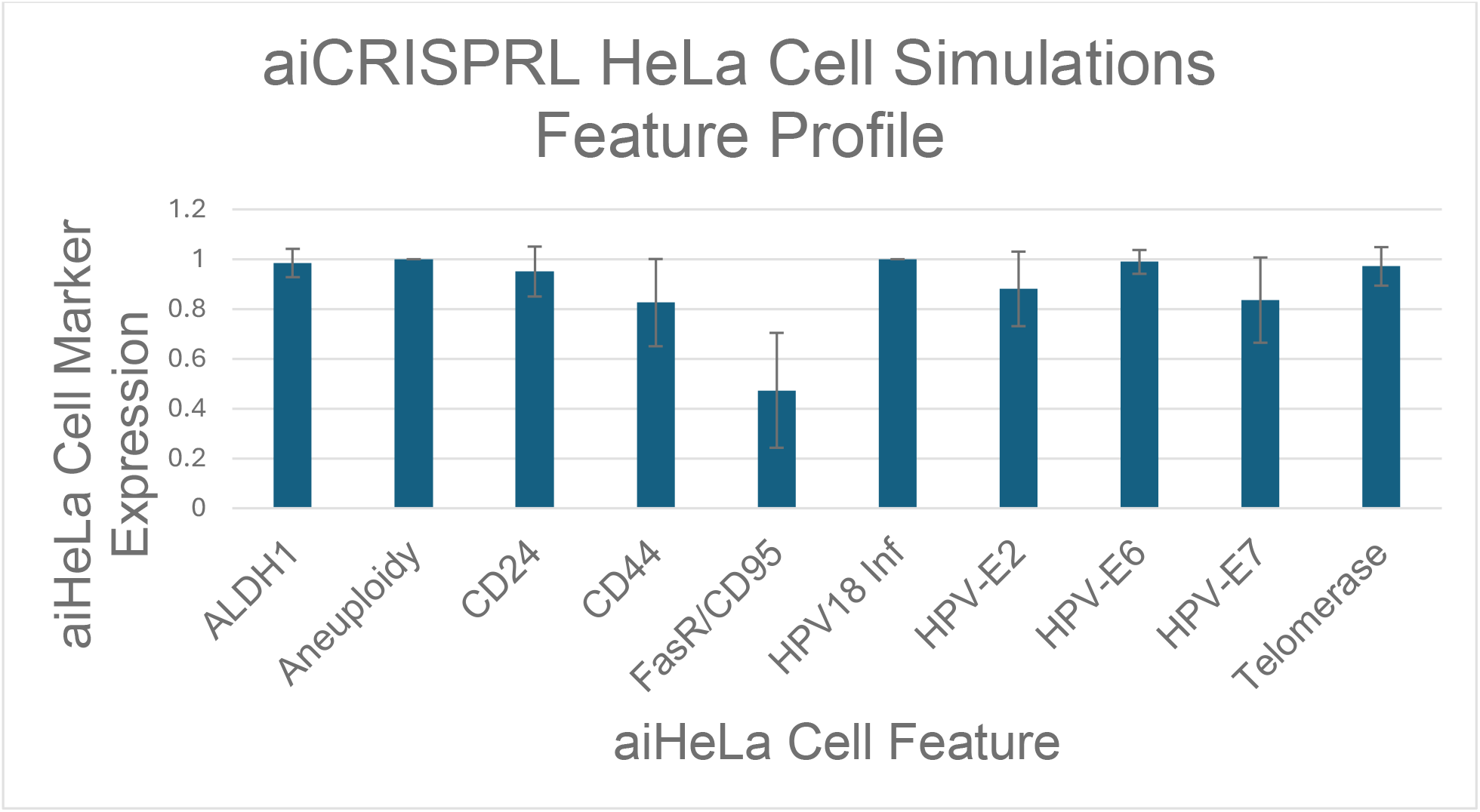
aiCRISPRL HeLa Cell simulations profile. aiHeLa Cell Marker expression. The vertical y-axis represents the semiquantitative levels of markers that are estimated by DeepNEU relative to an arbitrary baseline where 0 = none and +1 = maximum expression or presence. The horizontal x-axis represents the aiHeLa markers being simulated. Data represents means ± 95% confidence intervals. All p values are from the unbiased Binomial test. *p∼0.016.

### LOF Score Analysis

Analysis of the relationship between scaled MSH2 and the calculated LOF score revealed a strong, negative correlation with a Pearson r value of -0.884±0.035. A specific pathways analysis revealed a LOF for the extrinsic pathways of -0.819±0.042, and -0.965±0.020 for the intrinsic pathway.

Based on the sample size of 8 and a two-tailed critical r value of 0.765, the probability that this correlation occurred by chance alone is <0.01. Allowing for the scaling modifications to our MSH2 LOF score, the strong negative correlation is consistent with [27] in that more positive (or less negative in the case of aiCRISPRL) LOF scores imply deleterious mutations while more negative scores suggest neutral mutations. These results are presented in Table 4 and Figure 2.

**Table 4.** Legend: MSH2*= MSH2 gene expression scaled between -1 to +1 for input to the DeepNEU platform; 6-TG** = the predicted effect of 6-TG on aiHeLa apoptosis; Placebo*** = the predicted effect of Placebo on aiHeLa apoptosis; Ratio^#^= ratio of Placebo effect over 6-TG effect; LOF Score^##^= Log_2_(Ratio) as per [27]

| **Model-Average, N = 18** | **MSH2\*** | **6-TG\*\*** | **Placebo\*\*\*** | **Ratio<sup>#</sup>** | **LOF**<br>**Score<sup>##</sup>** |
| --- | --- | --- | --- | --- | --- |
| **aiHeLa_Cell (30/05/24)** | +1 | -0.746 | -0.615 | 0.824 | -0.279 |
| **aiHeLa_Cell (30/05/24)-AG11** | -0.400 | -0.643 | -0.603 | 0.938 | -0.093 |
| **aiHeLa_Cell (30/05/24)-AH07** | -0.300 | -0.641 | -0.623 | 0.972 | -0.041 |
| **aiHeLa_Cell (30/05/24)-DA02** | -0.904 | -0.344 | -0.389 | 1.131 | 0.177 |
| **aiHeLa_Cell (30/05/24)-DC08** | -0.932 | -0.358 | -0.368 | 1.028 | 0.040 |
| **aiHeLa_Cell (30/05/24)-KO** | -1 | -0.366 | -0.426 | 1.164 | 0.219 |
| **aiHeLa_Cell (30/05/24)-RD03** | -0.780 | -0.422 | -0.450 | 1.064 | 0.093 |
| **aiHeLa_Cell (30/05/24)-RH07** | -0.896 | -0.341 | -0.402 | 1.179 | 0.237 |

**Figure 2:**
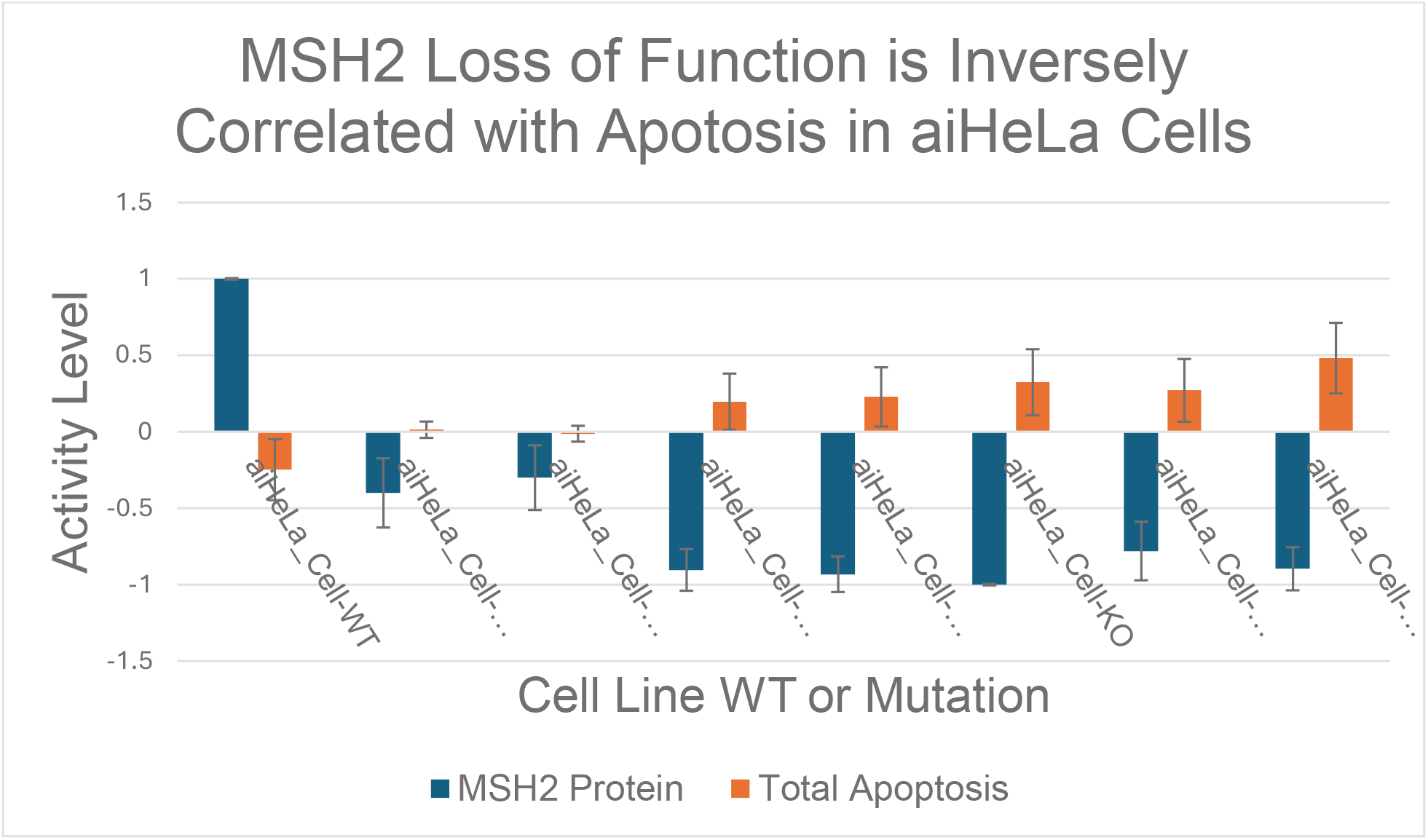
aiCRISPRL analysis of MSH2 Mutations in aiHeLa Cells. Analysis of the relationship between MSH2(scaled) and the calculated LOF score. Analysis of the relationship between MSH2(scaled) and the calculated LOF score, revealed a strong, negative correlation with a Pearson r value of -0.884±0.035, based on the sample size of 8 and a two-tailed critical r value of 0.765. Data represents means ± 95% confidence intervals, p<0.01.

Table 4 Pearson correlation (r) between MSH2 protein and LOF Score = -0.884±0.035

## DISCUSSION

Our previous successful gene editing applications of the DeepNEU platform are summarized in Table 1 above [5-7,10-21]. In the current project, we have extended those results by simulating HeLa cells and then introducing variable LOF mutations into a specific DNA damage repair gene (MSH2) in the aiHeLa cells. Based on a literature derived 10 element feature profile [22-26], our analysis of the data indicates that the aiCRISPRL application has successfully simulated a version of the original HeLa cell line (aiHeLa). Since 1951, the original HeLa cell genome has diverged to form numerous unique clones across the globe. These widespread changes in the HeLa genome have been the result of sequential infections from multiple bacteria, fungi, and viruses aided by more than 70 years of mutational pressures and rapid cell division. HeLa cells in culture have a doubling time of about 23 hours. As a result, currently available HeLa cells have little in common with the original cell line [29-32].

An important advantage of our DeepNEU/aiCRISPRL application for engineering aiHeLa cells allows us to avoid any unwanted external contamination from chemicals and infective organisms using a standardized culture medium. The only source of infection in these simulations comes from the simulated HPV-18 infection that generates E proteins 2, 5, 6, and 7. The impact of our early stopping regularization on system evolution is to reduce the impact of prolonged time on frequent DNA replication and rapid cell division. The use of a consensus approach to mutational profile was also designed to minimize the development of any mutations resulting from the combination of time, laboratory conditions, and contamination by multiple infecting organisms. Taken together these factors suggest that the simulations have produced a generic HPV-18 induced cervical cancer in a 31-year-old woman smoker and as a result, these simulations are likely to be more like the original HeLa cell type than current divergent clones.

Once the aiHeLa cell simulations were validated using the previously unseen published literature, the DeepNEU/aiCRISPRL platform was used to modify the aiHeLa cell MSH2 gene. This was conducted by introducing seven LOF mutations to produce a range of negative effects on MSH2 protein expression and then comparing these LOF mutations with the wildtype aiHeLa. The relationship between LOF mutations and the resulting MSH2 protein levels was scaled from the format in Table 3, to the DeepNEU input range where -1 = deletion of the MSH2 gene resulting in complete loss of protein and +1 = wild type gene status and normal levels of MSH2 protein.

The impact of these LOF mutations on aiHeLa cell DNA damage repair was evaluated by assessing the impact of 6-Thioguanine (6-TG) treatment on both WT and mutated aiHeLa cells as described in [27]. 6-TG is a thiopurine prodrug metabolized to its active form in the liver. The active form undergoes further metabolism to produce thioguanine nucleotides (6-TGNs) that can be incorporated into RNA and DNA synthesis as false purines resulting in potentially lethal DNA mutations. While the MMR process recognizes these mutations, it is unable to repair them resulting in replication arrest and apoptosis. Our analysis also confirmed that the effects of 6-TG induce apoptosis through the intrinsic and extrinsic pathways [35]. Additional cytotoxicity from 6-TGNs is the result of inhibiting the RAC1 protein that regulates the diverse downstream signals of the RAC1-VAV pathway in various cancer cells [34].

Consistent with the data from [27] we were able to confirm the effect of MSH2 protein loss on HeLa cell mortality as measured by the degree of apoptosis. When we analyzed the relationship between MSH2(scaled) and the calculated LOF score, a strong, negative correlation with a Pearson r value of -0.884±0.035 was revealed (N=8, critical r = 0.765, p<0.01). This finding confirms that the effectiveness of 6-TG to induce apoptosis in aiHeLa cells is dependent on a functioning MSH2 dependent MMR machinery. Importantly, aiCRISPRL editing of the MSH2 gene can accurately reproduce specific mutations and the LOF scores reported in [27]. In addition, this important relationship appears to be stronger for the aiCRISPRL editing case (Pearson r = - 0.884±0.035 vs 0.770).

### Conclusions and Future Considerations

In this study, we successfully employed the DeepNEU platform to simulate aiHeLa cells that accurately resembled the original and uncorrupted immortalized HeLa cell line. We then evaluated the DeepNEU derived aiCRISPRL like capabilities to introduce a series of specific LOF mutations into the MSH2 gene. We chose to study the MSH2 gene because it is a critical component of the MMR DNA repair pathway and it could be evaluated by treating the affected HeLa cells with the cytotoxic prodrug, 6 Thioguanine (6-TG), and observing the degree of resulting apoptosis. The severity of the loss of function mutations in MSH2 was estimated from the LOF score. This score was calculated in a manner like that reported in [27] and produced data confirming a highly significant inverse correlation between MSH2 protein levels and aiHeLa cell apoptosis. This methodology has permitted the direct comparison of CRISPR-Cas9 and aiCRISPRL technologies for introducing specific MSH2 mutations into the HeLa/aiHeLa genome and while the technologies are different, they are directly comparable. Furthermore, like CRISPR-Cas9, aiCRISPRL can be used to create single, multiple, and sequential (LOF and GOF) mutations (see Table 1).

CRISPR-Cas9 is a mature and impressive in vitro gene-editing technology with an improving success rate while aiCRISPRL is a specific application of an evolving AI platform technology that can be quickly and reliably deployed. Like other AI simulation technologies, DeepNEU requires substantial amounts of validated data from multiple sources. The current DeepNEU network is composed of ∼5.7X10^7 artificial neurons and the database contains relationship data for ∼34% of the human genome. Our goal is to obtain relationship data for ∼99% of the human genome over the next 2-3 years.

Finally, we have evidence that the DeepNEU platform has recently entered the domain of Wise Learning (WL) as it relates to health care. The WL process defined by Groumpos in 2016, represents the next evolutionary step in AI that combines Fuzzy Cognitive Map simulations, with data from multiple experts and a Generic Decision-Making System (DMS). The WL process should also explore available learning algorithms including Deep Learning (DL) methods when available [33]. While the DeepNEU platform continues to evolve, as of this writing, the current version (8.4) meets all these Wise Learning stated criteria.

## Acknowledgement

The author thanks Dr Sally Ezra (PhD) for her expert and thoughtful input to an early draft of the current manuscript.

